# On a simple general principle of brain organization

**DOI:** 10.1101/771535

**Authors:** Jose L. Perez Velazquez, Diego M. Mateos, Ramon Guevara Erra

## Abstract

A possible framework to characterise nervous system dynamics and its organization in conscious and unconscious states is introduced, derived from a high level perspective on the coordinated activity of brain cell ensembles. Some questions are best addressable in a global framework and here we build on past observations about the structure of configurations of brain networks in conscious and unconscious states and about neurophysiological results. Aiming to bind some results together into some sort of coherence with a central theme, the scenario that emerges underscores the crucial importance of the creation and dissipation of energy gradients in brain cellular ensembles resulting in maximisation of the configurations in the functional connectivity among those networks that favour conscious awareness and healthy conditions. These considerations are then applied to indicate approaches that can be used to improve neuropathological syndromes.

## 1 Introduction A possible framework to characterise nervous system dynamics and its organization

The emergence of consciousness and self-awareness from nervous system tissue –connected in turn to other organs and immersed in an environment– is a topic of considerable interest for which no clear answer has been found. Consciousness as an emergent property of the brain is a notion that many scholars support. However, this does not provide much information as it does not shed light into mechanisms through which conscious awareness proceeds. Considering consciousness as the psychological state of conscious awareness, there are aspects of consciousness which can be investigated in depth –like memory formation, volition, self-awareness etc.– and can provide insight into the search for a general principle of nervous system organization and its associated dynamics that results in conscious awareness and in unconscious states, both normal and pathological. Following J. Williard Gibbs advice that “One of the principal objects of theoretical research in any department of knowledge is to find the point of view from which the subject appears in its greatest simplicity” (J.W. Gibbs in a letter to the American Academy of Arts and Sciences, 1881), we turn to one aspect that we, and many other investigators, posit is fundamental for the development of conscious awareness: the transient establishment of connections among brain cell networks –which we will assume are neuronal networks even though the activity of glial cells have also important consequences for the contacts among neurons. Focusing on the distributed interactions that underlie the system’s collective behaviour could be a useful start in the search for simplification. This aspect will be investigated from a high level perspective, being the main purpose the attempt to find a principle of brain organization and associated dynamics that encapsulates the emergence of conscious awareness and the deviant progression to neuropathological states.

In the search for a simple unified conceptual framework to describe the organization of brain dynamics we aim to bind some results together into some sort of coherence with a central theme. We follow the thermodynamic approach: search for a state functional that reflects the nature of the states attained by the system and that is influenced by certain observables. Two main considerations are in order to determine what constitutes the states of the brain, what observables can be measured and what level of description is the most relevant for the purposes of characterising brain and behaviour.

First consideration, brain activity is normally described from EEG, MEG or functional neuroimaging as a superposition of dynamics at different time scales; considering what is known about how the nervous system operates during cognitive states, it is conceivable to infer that the “nature of states” consists of patterns of neuronal coordinated activity –that is, correlations of activity. Hence, coherence or synchrony could be a fundamental observable. It is already accepted that the evaluation of neural synchrony constitutes an appropriate metric to characterise nervous system dynamics; as stated by Mora & Bialek (2011) “correlations strongly determine the global state”. Furthermore, temporal coordination of neuronal activity is essential for cognition in general [1]. There are scores of studies suggesting the fundamental importance of correlated activity among neurons for sensorimotor integration and in general to process information; just two examples: in moths the pattern of neural ensemble synchrony is best correlated with behaviour [2], and odour representation in zebrafish is mediated by the coordinated responses in neural ensembles rather than by other measures like absolute neuronal activity [3]. On the whole it can be asserted that the foundation of neural information processing lies in the interaction between cells, each cell’s activity is meaningful only with respect to other cell’s actions. Thus, synchrony (or correlations) of cellular activity can be the observable that influences the patterns of organised activity that constitute the brain states. Hence, it is conceivable to propose that, to follow the abovementioned thermodynamic approach of finding a state functional that is influenced by observables and that reflects the nature of the states attained by the system, the nature of states are patterns of coordinated activity that can be observed as correlations/synchronization of activity in cell ensembles. Measures of synchrony, then, become our relevant observable.

A second consideration refers to equilibrium, as it is known that life phenomena are far from equilibrium. Because there are advantages in the analysis of complex systems when notions related to equilibrium are used, the question becomes whether these concepts can be applied to the nervous system. Equilibrium holds when the macroscopic parameters of a system are approximately constant, and because “the concept of equilibrium state is intimately connected with that of observation time” [4], then –just like the Markov process concept– there is a time scale (and a localised region) at which a system can be found very close to equilibrium, and in fact there exists the concept of local equilibrium [5]. The question of steady states being at equilibrium or not may be a matter of scale of observation. It is thus of interest that when a system is observed at coarse-grained scales –which is a very relevant scale for neuroscience to understand the relation between brain and behaviour, that is, the mesoscopic level [6] which throughout the following text will be taken as the synchronization among cellular networks– far from equilibrium systems recover thermodynamic equilibrium properties at these coarse-grained scales [7]. In addition, it has been argued that the EEG exhibits near equilibrium dynamics at macroscopic scales, despite the nonlinearities at the microscopic level [8]. Hence, it is conceivable to apply some equilibrium thermodynamic notions at the macro/mesoscale level of brain activity. In fact, the field of cognitive thermodynamics may have already been created [9], and attempts to characterise cognition from a thermodynamic perspective have been advanced [10] with some early pioneering studies already proposing thermodynamical models of brain activity that, perhaps due to the lack of precise data of nervous system’s observables in those early times, were somewhat vague but illuminating [11]. Let us digress briefly to note that we use the term “network” quite often in our narrative, and it is not easy to precisely define what constitutes a brain cell network in a manner that satisfies everybody. In reality networks are sort of abstractions, just like “brain state”, but useful for neuroscientists to communicate. From our perspective we will consider networks the transient functional groupings of active cells, using [12] definition: “gradual clustering according to similar activity profile”.

To sum up, it can be conceived that for our purposes of finding general principles of brain organization the adequate level of description is associated with the mesoscopic correlations of activity –e.g., coherence or synchrony between brain cell networks– which will allow for a careful application of thermodynamic concepts. Furthermore, it has to be considered that cell networks coordinating their activity act as dissipative structures (just like any other biological phenomenon), giving rise to energy gradients such that “mental things happen”. A primordial consequence of the neuronal energy moving down gradients is the fluctuations in the correlated actions of cell networks, or in other words the instability –or metastability [13]– of brain states: very stable states are normally pathological, like status epilepticus or coma. In general, energy moving down gradients constitutes a process involving determinism (cooperation/competition) and stochasticity, which results in pattern formation in natural phenomena [14]; this interrelationship between determinism and stochasticity in pattern formation can be clearly experienced in the case of dynamic random fractals [15]. This indicates that a probabilistic framework interpretation of brain activity may be very useful. The following scheme (figure 1) for the search of a general principle of brain function emerges.

**Figure 1:**
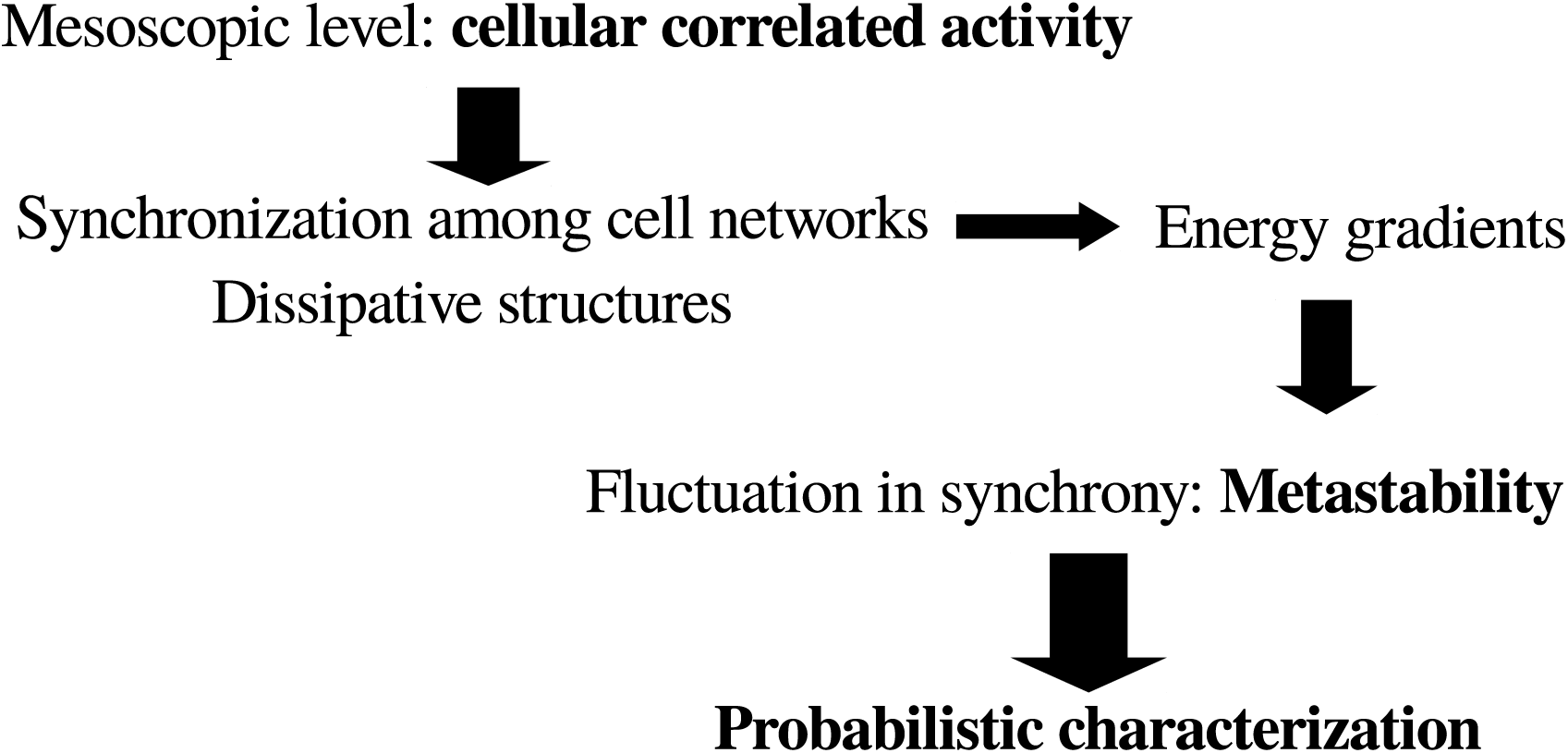
General scheme towards a possible framework to characterise brain dynamics in the search for a basic general principle of brain and behaviour. See text for details.

The text is organised according to the following logic. Based on previous experimental observations on the entropy associated with the number of configurations of connected brain networks, some considerations on the free energy available in conscious and unconscious states are described. Then energy dissipation will be considered, as the lower free energy in conscious states may indicate higher dissipation, and we will see that the neurophysiological tendency towards greater dissipation in healthy and conscious states emerges. The neurophysiological importance of dissipation will be considered next, and in section 5 some approaches that can be used to improve neuropathological syndromes are described, based on the previous sections on the basic thermodynamics of cognition. Section 6 describes a probabilistic characterization of brain dynamics after the observation that the dissipation of energy gradients is associated with large number of configurations of neural synchrony patterns, in an attempt to find clues about the dynamical evolution of brain microstates that compose the macrostates with different number of connections. Finally, section 7 presents some thoughts of the emergence of cognition derived from the notions of dissipation and compartmentalization.

## 2 Thermodynamic considerations regarding brain dynamics in conscious and unconscious states

The aforementioned comments on the applicability of equilibrium concepts to the mesoscale level allow for the use of notions like free energy, using the standard Gibbs free energy equation *G* = *E*−*TS* with *E* the internal energy, *T* the temperature and *S* the entropy (there is also the analogous Helmholtz equation *F* = *U* −*TS*, but in what follows the standard Gibbs expression will be used). In general, for nonequilibrium systems, the terms in this formula are not the exact terms used in classical thermodynamics of equilibrium systems. For non-equilibrium one has to consider functionals of a similar form associated with a certain probability distribution *P*, *G*[*P*] = *E*[*P*] −*TS*[*P*] where *E* is some kind of internal energy, *T* is a noise term representing fluctuations, and *S* is an information measure like Shannon entropy, which is formally equivalent to the Gibbs entropy [16].

There has been a recent interest in the relevance of a free energy perspective on cognition [17]. It is known that nature follows the path towards a decrease in free energy, the dissolution of energy gradients. It is crucial to note too that it is not absolute energy (as energy is hard to define) *but the differences in energy that have physical meaning* [18]. At or near equilibrium the distribution of stationary states corresponds to those that minimise *G*. Along these lines, regarding the nervous system, it has been said that “consciousness integrates sensory and other inputs [….] to consume energy gradients more effectively than by unconscious deeds” [19]. Hence, is conscious awareness associated with lower free energy in the brain?

A tendency in the evolution of brain free energy (*G*) is all that can be uncovered since a precise estimation of *G* is very difficult because the terms cannot be accurately estimated, unlike in chemical reactions. The term *T* can be considered noise, *E* the brain internal energy and *S* the entropy associated with the number of configurations of connections among neuronal networks. Other entropies associated with other brain phenomena can possibly be chosen, and the choice of the observable to compute entropy may of course change some results, a similar situation as occurs with the application of several complexity measures to neurophysiological data that give different results depending on what the complexity is measuring [20]. Since we are considering the mesoscale level of cellular collective activity, the entropy associated with connectivity patterns is the one considered here. While this entropy can be accurately calculated (see below), it is unfeasible to precisely compute the noise term, *T*, even though considering it as fluctuations it could be in principle possible to calculate fluctuations in specific observables. Since *S* will be, in what follows, the entropy associated with connectivity patterns derived from neural synchronization analysis, then it is logical to conceptualise here the noise term as fluctuations in synchrony; in previous work, synchronization among brain signals has been used and the fluctuations in synchrony evaluated [21, 22]. Consequently, *T* can be considered fluctuations in the functional connections among cell networks. The rationale for this particular notion of brain noise is that cellular correlated activity, as previously remarked, is fundamental for appropriate nervous system function. There are other observables from which noise can be estimated but to be consistent with past studies that will be used in what follows, where the entropy was quantified from synchrony patterns, it makes sense to describe the noise term as fluctuations in synchronization. As well, the internal brain energy *E* is equally impossible to estimate. What is known is that brain energy –normally assessed taking as proxy cerebral blood flow, oxygen consumption or glucose uptake –is basically constant regardless of mental effort or state. It tends to diminish in pathological conditions like vegetative and minimally conscious states or coma [23] and it tends to increase in epileptiform activity although this activation is restricted to brain areas involved in the seizures [24]; during sleep the global change in brain energy is unclear because there are various brain regions that are metabolically more active and others less active [25], and even a surge in ATP levels in the initial hours of sleep has been reported [26], all of this making it difficult to ascertain whether there is a net change in brain energy during sleep even though it is agreed that there is a tendency towards lower metabolism [27]. These comments reflect the difficulty in assessing brain energy, and again only tendencies are found, namely, that there is hypometabolism in coma and sleep, and hypermetabolism in seizures.

On the other hand, it has been proposed that energy for brain macrostates could be derived from a measured probability distribution function (pdf), in that an energy attribute can be assigned to every state of the system from the pdf of microstates [28]. This comes from the celebrated Boltzmann expression *P* = *e*^(−*E/kT*)^, so a brain macrostate energy is *E* = −*T* Σ_*m*_ ln(*p*_*m*_) assuming that *k* = 1 and the summation is over all m microstates (with probabilities *p*_*m*_) making up the macrostate. Can this trick be used here?

In previous studies we obtained the pdf of connected signals in different conscious states, the microstates that were used to calculate the entropy of the configurations of connections [29], thus in principle those pdf could be used to estimate each *p*_*m*_ and assign an energy to the corresponding macrostate, resulting in an expression for the free energy of each cognitive/brain macrostate *G* = −*T* [Σ_*m*_ ln(*p*_*m*_) + *S*] where all terms could be quantified including the noise as the aforementioned quantification of the fluctuations in synchrony. Energy in probabilistic terms with possible applications to neuroscience has been advanced as well by others [19]. But there are two inconveniences with this approach. The first is that to compute the entropy *S* it was assumed equiprobability of the microstates making up a macrostate (details in [29]), therefore it would be erroneous to mix in the same expression the notions of equiprobability of microstates and that of the probability depending on the energy of each microstate. A second problem derives from the interpretation of this sort of abstract energy; if we were allowed to estimate the brain energy as aforesaid, the energy for the highly connected states like seizures or coma would be high because the probability of the macrostate seizure or coma is low, but as described above there is a tendency towards hypometabolism in coma, hence a contradiction appears. The state of lowest energy would be the fully awake state as it is the more probable, but again an inconsistency occurs because, as aforementioned, there is a tendency to more metabolism during awake states than during sleep. Therefore, the neurophysiological interpretation of this abstract energy derived from the pdf is not trivial. Nevertheless, and to close this digression on brain energy, from a thermodynamic perspective we can talk about metabolic or any other type of energy, and work with it, as already anticipated by pioneering studies: “Although the inflow of free energy to the brain may appear as electromagnetic signals or as chemical free energy, the thermodynamic model need not, in view of their physical equivalence, distinguish between them” [11].

Still, after considering all these limitations in the application of the expression for free energy to the brain in different states, some tendencies in brain free energy *G* can be inspected. In previous work we used invasive and non-invasive electrophysiological brain recordings in different states of consciousness to determine the entropy *S* associated with the number of possible configurations of pairwise connections; that is, the phase synchrony between pairs of brain signals was evaluated as regularly done using the analytic signal concept and a threshold value of the synchronization index was used to determine when two signals were “connected” (details in [29, 30]). The signals are assumed to correspond to the activity of a network of cells located in the neighbourhood of each electrode/sensor, hence in what follows we shall use the term cell network as synonymous of signals. Each state of consciousness studied (wakefulness, sleep, coma and seizures) was considered a macrostate of brain synchronization, composed of a number of microstates which are the several possible configurations of the connected signals. For the current purposes of applying these considerations to inspect the trends in the evolution of free energy *G*, in the expression *G* = *E* − *TS* for each macrostate, *S* will be the abovementioned entropy, *T* we shall assume that is analogous to noise or fluctuations in synchronization that can be evaluated as *dN*_*i*_*/dt* with *N*_*i*_ being the pairwise connected networks, that is, *T* is the variation in time of the pairwise connected brain networks; and *E* will be qualitatively taken as the trends in brain energy associated with different cognitive states mentioned three paragraphs above.

The results obtained in the aforementioned studies showed that the entropy associated with conscious awareness was higher than that of unconscious states (Figure 2 summarises the main findings that will be used in the following arguments). It is fair to mention that since the *S* calculated was based on a measure of phase synchrony, it varied depending on the frequency used to evaluate the synchronization between the signals. Low frequencies (*<* 10 Hz) differentiated best conscious versus unconscious states; the explanation for this observation was presented in [30]; here, suffice to say that there is evidence supporting the fundamental importance of the slow brain rhythms for brain information processing and how higher frequencies “ride on” the slow waves. The communication among neurons via nested rhythms is a known phenomenon [31] and low frequencies serve as temporal reference for information transfer at other, higher frequencies. As well, entropy applied to another feature of the brain signals (spectral entropy) was found lower in unconscious states (acute and chronic disorders of consciousness) as compared to healthy individuals [32].

**Figure 2:**
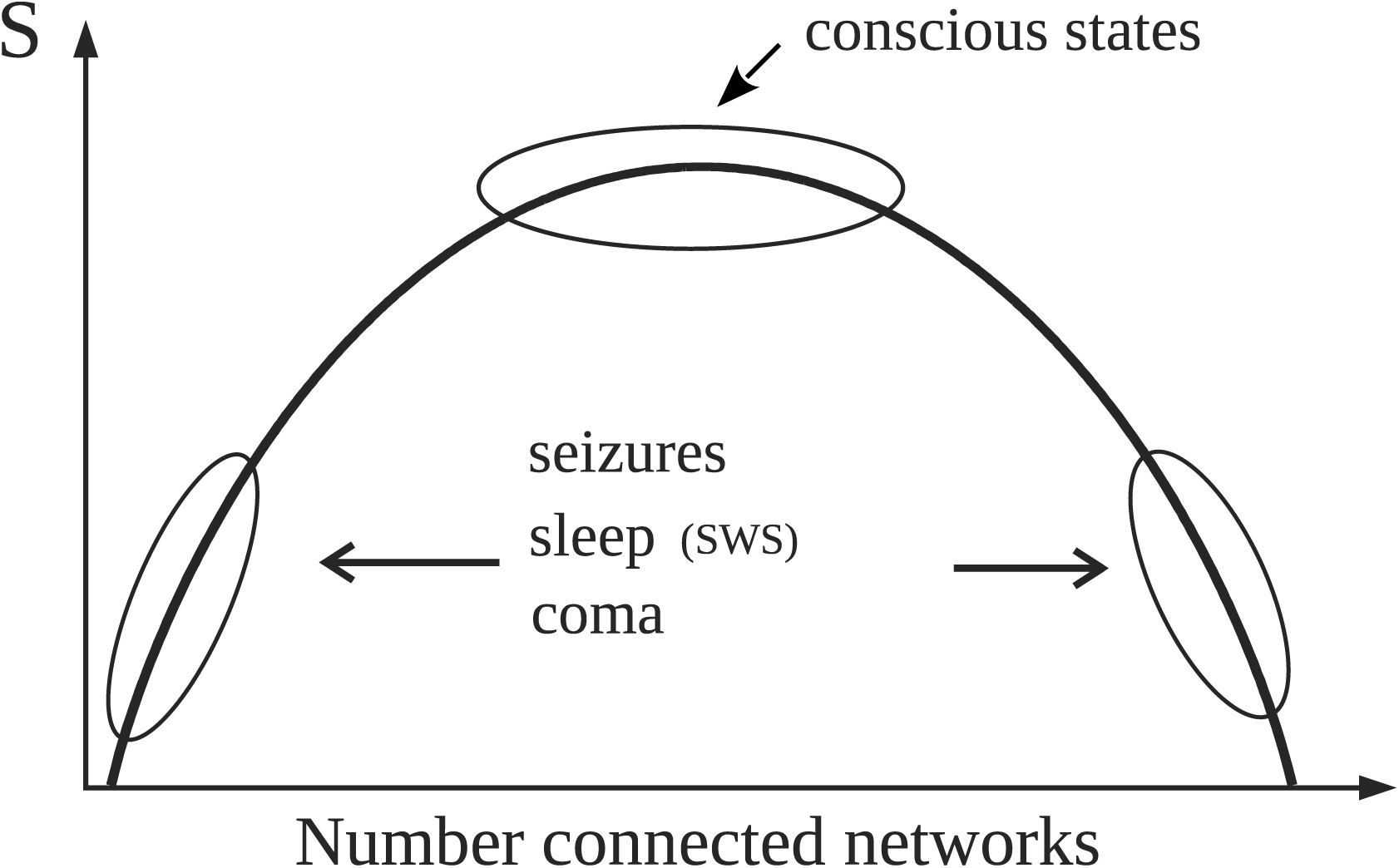
Simple schematic depicting the main results reported in [29] that are used in the text (please see technical details in the original publication). Normal alertness resides at the top of the curve representing the entropy (*S*) associated with the number of configurations of connected brain cell networks. The top of the curve with high *S* represents large number of configurations (microstates) of network connections which provides the variability in brain activity needed for normal cognition. Pathological or unconscious states like slow wave sleep (SWS) are located away from the top, thus characterised by either large or small number of “connected” networks therefore exhibiting lower number of microstates, hence lower entropy. As detailed in the text, for constant noise (*T*) and energy (*E*), *G* will be then lower near the top of the curve because of the high values of *S*

Hence, due to the higher value of entropy in alert conditions, for constant *E* and *T* the free energy *G* should thus be lower in conscious states, which would indicate that the energy gradients are better consumed (that is, dissipated) during conscious awareness, as proposed by A. Annila (2016). Now, neither *E* nor *T* remain constant for different states of consciousness, as previously mentioned. In conscious awareness characteristic of wakefulness where the internal energy and the noise (fluctuations in synchrony) are higher than in unconscious states [21, 22], the entropy should increase in order to follow the natural tendency to decrease *G*, which is what was found in our studies as mentioned above: high *S* values in alert states (Figure 2). On the other hand, during unconscious physiological – that is, normal – states (sleep) with lower noise, entropy does not need to be prevalent and also considering that there is hypometabolism during sleep, then *S* can be lower than during wakefulness to still have a diminished free energy; and indeed lower *S* associated with slow wave sleep (SWS) was found (figure 2 in [29], schematised in the current Figure 2). In unconscious pathological states (coma, seizures), where lower entropy was found as well, the evolution of *G* will depend on the energy and the noise. In coma both brain energy and noise – again, noise as fluctuations in neural synchrony [22] – are low, therefore *G* will be basically equal to the internal brain energy which may be low due to the pathological hypometabolic state; whereas during seizures the energy increases while the noise decreases [21], thus *G* will tend to be larger in the pathological state of epileptic seizures. In general, waking conscious states minimise *G* better than unconscious states.

To maintain healthy brain states then is not about the total amount of energy in the brain, as there seems to be no clear relation between the energy used and the degree of conscious awareness (e.g., large energy in seizures with loss of consciousness), but rather in how the energy is organised, namely, how the brain cell networks coordinate their activity and distribute the energy to process sensorimotor information. Same can be said about cellular activity, which of course runs in parallel to energy consumption: High brain activity is not consciousness; rather, it is a property that provides necessary but not sufficient support of the conscious state [33]. The functional importance of the organization of energetic processing in the brain has been proposed by some authors [34], which is line with older proposals that postulated that the flow of energy through a system acts to organise the system [35] and that biological information is, in the final analysis, the energy flow [36] as metabolic pathways offer channels for energy to flow and cellular communication provides further conduits for that flow. Not only in the biological world but also in the inorganic, channels exist so that energy flows and creates patterns [14]. Therefore putting together these considerations about brain energy and neuronal activity seems to lead to the notion that the crucial aspect for proper conscious awareness is that brain cellular functional connectivity and activity has to be variable enough –provided by the creation and dissipation of energy gradients– and well organised so that the integration and segregation of information, two fundamental aspects for proper sensorimotor processing, can occur. The view that consciousness relies on large–scale neuronal communication is a common point of several cognitive theories [37].

## 3 The neurophysiology of dissipation

The question of how brain free energy changes with healthy or pathological brain states can be inspected from another perspective. It has been proposed that brain dissipates energy in order to process information [34, 38]. As a dissipative structure, the nervous system should then dissipate energy efficiently to function properly, and the question becomes whether there is a relation of dissipation with healthy and pathological conditions. Building on the studies of [39] the energy dissipated during avalanches, some considerations can be obtained about brain dynamics especially reflecting on energy dissipated in the process of synchronization of brain cell networks and how it changes with neuropathology. The authors propose that the size of an avalanche is proportional to the instantaneous energy dissipation rate. In neuroscience, neural avalanches have been considered in the literature, normally as bursts of activity in neuronal networks (Beggs and Plenz, 2004). Those bursts of activity occur due to neurons receiving synchronous inputs from other connected cells, thus, in the final analysis the bursts represent manifestations of synchronization of cellular activity, which naturally correlate with the amplitude of the extracellular field potentials: large amplitudes represent more synchronous cellular activity; hence, synchrony and magnitude of local field potentials are very much related.

Neural synchrony in neuroscience is evaluated using certain indices. Let us use one originally described by Mormann et al. (2000), a mean phase coherence statistic which is a measure of phase locking and defined as *R* = time |⟨*e*^*i*Δ*ϕ*^⟩| where Δ*ϕ* is the phase difference between two signals. The magnitude of *R* normally evaluated within a certain window of a few milliseconds to 1 second describes how synchronous two signals are, signals which from now on will be referred to as networks because the signals represent activity of many neurons located nearby the recording sensor/electrode. The larger the magnitude of *R* the more synchronous two networks are, which indicates that more neurons are becoming entrained in both networks (recall we are talking about signals representing collective activity and not two individual neurons connected, because in the latter case there could be changes in R without changing neuronal firing activity); here we assume, for the sake of simplicity, that the cell networks are directly connected there could be enhanced synchrony as well if both brain regions are receiving common input from a third one, but in the end the reasoning, while more elaborated, results in similar final arguments. Therefore, it is conceivable that the magnitude of *R* represents the size of the neural avalanche. Following [39]considerations, we have 𝒮_*z*_ ∼ ∫ *F* (*t*)*dt* where 𝒮_*z*_ is the size of the avalanche and *F*(*t*) the energy dissipation rate. If we then assume that 𝒮_*z*_ –in this case the clusters of connected cells is equivalent to the synchronization index *R*, it leads to the following relation *F* ∼ *dR/dt*

From this simple relation between the dissipation rate (*F*) and the time evolution of synchrony (*R*) some indications emerge about the dissipation of energy in brain activity. Energy is dissipated as more neurons become connected, thus when *dR/dt >* 0, and if the connections remain constant then *F* = 0, no dissipation occurs. Obviously, this reasoning is only applicable to the meso or macroscale, but not to two individual neurons becoming more or less synchronous, as there is no cluster in the case of two coupled neurons. From the above expression, it is then conceivable that the more abrupt the changes in *R*, the larger the dissipation; a sign of the tendency to dissipate energy can thus be seen by inspection of the evolution of the phase synchrony index R: the usual time series of *R* displays a jagged, spiky evolution (Figure 3). It reflects the tendency of cell networks to synchronize and desynchronize under the influence of perturbations, either external or from internal stochastic fluctuations that desynchronize cell ensembles. In pathological states like seizures there is an almost constant high value of *R* with lower fluctuations in synchrony [21] –as depicted in Figure 3– and in coma after traumatic brain injury a similar less variability in *R* was found [22], hence in these conditions less dissipation is expected according to the present argument. Therefore similar conclusions as those presented in the previous section about brain free energy are reached after these considerations. To say that when the changes in synchrony between neural networks are rapid the larger the dissipation becomes is another way to emphasise the importance of the variability in synchrony patterns, which is already becoming a common conclusion throughout this text.

**Figure 3:**
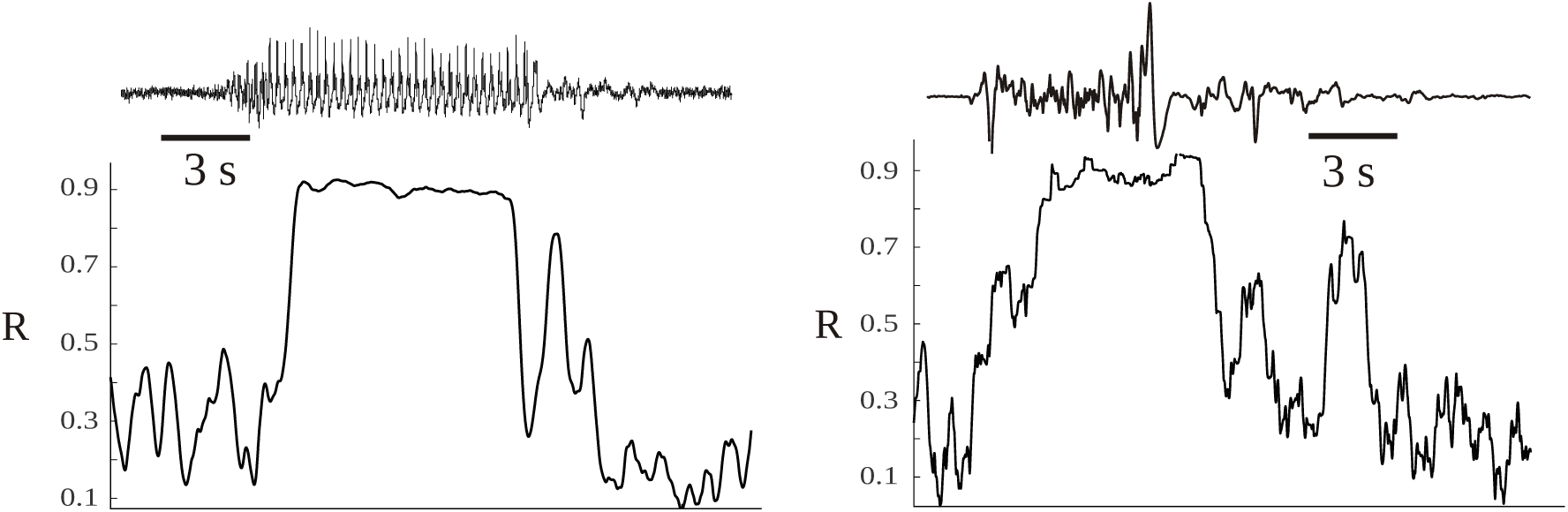
Time evolution of the synchrony index R between two magnetoencephalographic (MEG) signals taken in two epileptic patients during seizures. The patients were two of those described in [40], having two different epilepsies. The left hand side corresponds to an absence seizure (one MEG channel shown above, the ictus occurs during the high amplitude signal) and the other to a tonic seizure. During the seizures the synchronization index is high and shows less variability, losing the characteristic spiking as in moments without the ictus.

Another indication of dissipation can be obtained from the basic formula of macroscopic thermodynamics of irreversible process [5] *dS/dt* = *J*_*i*_*X*_*i*_ where *J* are the rates of the processes involved and *X* the generalised forces. This expression –in chemistry where it is commonly applied also appears under this fashion (basically using the chain rule) *dS/dt* = Σ_*i*_(*dS/dN*_*i*_)(*dN*_*i*_*/dt*) where *N*_*i*_ are molecules involved in the chemical reaction. The extension of this expression from chemistry to neuroscience can be envisaged if we consider not molecules but neuronal networks, specifically pairwise connections of networks, such that Ni are pairwise connected nets. Therefore, the term *dN*_*i*_*/dt* represents fluctuations in synchronization as discussed in the previous section, that is, the variation of the pairwise connected brain networks. This interpretation is along the line of dissipation viewed in terms of the emergence and vanishing of the coherence (connections) between cells (Torday and Miller, 2016). More fluctuations –larger number of configurations and variability of connected networks– were found during conscious awareness as opposed to unconscious states [29, 30], therefore suggesting that *dS/dt* is larger during wakefulness (as was concluded in the previous analyisis): the more fluctuations the larger the dissipation. In situations when fluctuations in synchrony are very low, like during seizures or coma, *dS/dt* should diminish. Once again the neurophysiological tendency towards greater dissipation in healthy and conscious states emerges.

## 4 Neurophysiological importance of dissipation

Why could dissipation be important for consciousness and healthy brains? In general, dissipation of energy is at the basis of pattern formation [5], and in the case of brains the patterns of organised cellular activity is what determines behaviour. Differences in energy –that is, gradients– is what makes things happen. In the case of nervous systems, those gradients are associated with communication among cell ensembles that carry out the fundamental aspects of sensorimotor processing, executing adaptive behaviours. Thus, at one level of description the coordinated cellular activity can be studied using the typical methods of coherence, phase synchrony, mutual information and the like, and at another level these collective patterns of activity can be viewed from a more abstract perspective employing the thermodynamical concepts of energy dissipation and entropy production.

It is gradients of energy that are in fact recorded in any typical neurophysiological signal, as shown in the two examples in Figure 4. These are bipolar voltage time series of about 1 second duration recorded from the interior of the brain of a rat, and the signal waveforms are due to differences in voltage; without these gradients, only flat lines would be recorded. In one recording a large amplitude rhythmic waveform is shown, and at this moment the rat was suffering an epileptic seizure; the other signal was taken at a time when he was not seizing. Notice the difference in the amplitude, the large amplitude during the seizure is due to many neurons firing in synchrony, whereas the healthy signal depicts the typical low amplitude high frequencies (gamma range) associated with wakefulness. For the sake of simplicity, let us imagine that the bipolar recordings represent the differences in voltage between to neural networks. The many tiny spikes that are seen in the low amplitude trace during normal behaviour suggest that there are several subnetworks within the two networks that are exchanging information, many gradients of energy (potential differences) allow many neurons to participate in several connection patterns, and the amplitude is small because there are few cells in each subnetwork; but the main point is that there are many subnetworks that communicate with one another thus each of the small spikes can be thought of as being created by the communication between two small cell populations, and because there are many small peaks this suggests there is great variability in the configurations of the connections among the neurons in both networks. The net result is the emergence of gamma frequencies, so much talked about with regards to cognition. Gamma frequencies reflect the emergence and dissolution of communication among small neuronal ensembles, fluctuations which provide variability needed in brain dynamics to properly process information leading to adaptive behaviour. On the other hand, the large amplitude signal during the seizure with less high frequencies indicates that many or perhaps almost all neurons in the two networks are involved in the communication, and therefore there is not much flexibility in terms of the possible configuration patterns that can be created: the potential differences are large because so many cells are synchronously active but there are very few gradients as compared with the previous condition, in fact, there could be only one gradient if just one perfect sinusoidal-like waveform were recorded (but this never happens in real life, only in computer simulations). In this manner, the origin and function of the gamma rhythms lose part of the mystery: the prevalence of gamma frequencies during wakefulness is just a matter of probabilities, it is probable that the great variability in the firing patterns and configurations of neuronal connections during cognition generates many voltage gradients. There have been proposals relating gamma frequencies to the infrastructure of the brain neuronal operations that balance excitation with inhibition when the brain is engaged functionally with the environment [42].

**Figure 4:**
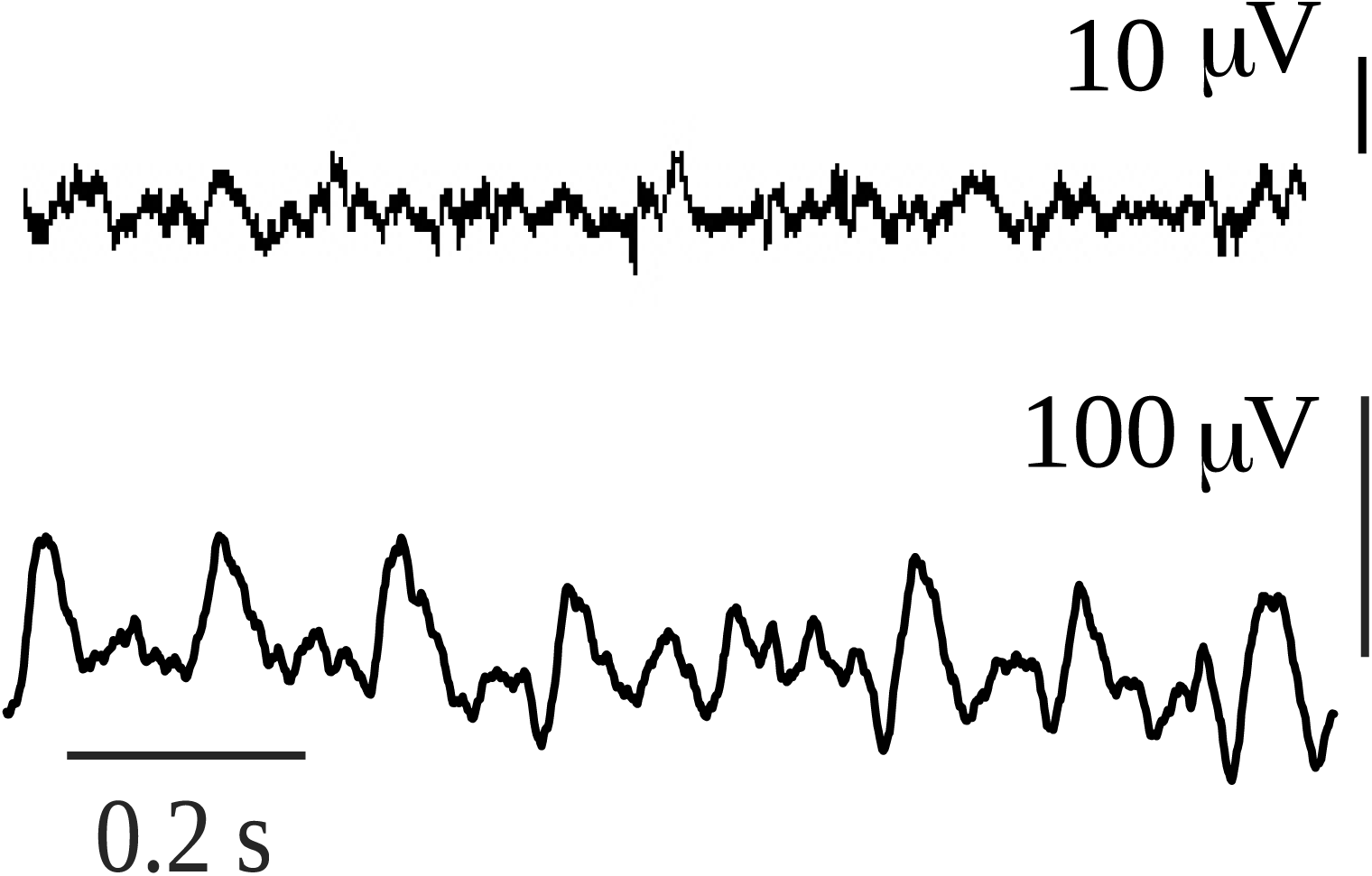
Intracerebral recordings of 1 sec duration in a rat that had spontaneous absence seizures. The first recording was taken during normal exploratory behaviour of the rat, while the second corresponds to an absence seizure. The animal model and the intracerebral recordings of this type of absence seizures have been reported in numerous publications (e.g. [41]).

From an evolutionary point of view Bruner et al. [43] described that the phylogenetic evolution of the human genus was associated with the increase of the functional and structural brain complexity. This is due to the necessity to handle the information (internal and external) in the most efficient way. As we discussed before, the brain processing of information has associated energy dissipation. Therefore, we could conclude that as the brain evolved over time the energy dissipation increased.

In conclusion, the creation and dissipation of energy gradients is associated with brain conscious awareness and healthy conditions, which may indicate approaches that can be used to improve neuropathological syndromes.

## 5 Improving healthy brain dynamics

Based on the observations that conscious awareness and healthy brain states are characterised by higher entropy, decreased free energy and increased dissipation, one adventurous speculation is offered as to how to improve the health of the brain in pathological states, especially of low entropy. Based on the previous results, increases in the dissipation –that is, reductions of free energy– may provide useful to correct some pathological states. The basic aforementioned formula *G* = *E* − *TS* provides some hints. Considering that the brain internal energy is high and almost constant during healthy states, then perhaps it should not be lowered, and in any event it is very difficult to manipulate brain’s energy. The safest method to decrease the free energy would thus be to increase the noise. This can be done using neurostimulation; for instance, deep brain stimulation (DBS). It is thus of interest that DBS has been used in a patient in minimally conscious state who was partially recovered by stimulation of the intralaminar thalamic nuclei [44]. The neurophysiological basis of the effect of this stimulation can be understood by considering the net effect of increasing excitability in the thalamus: because the intralaminar nuclei project to almost all brain cortical areas, the stimulation causes an overall increase in cortical excitability, that is, enhances internal brain noise. In another pathology, epilepsy, noisy or asynchronous stimuli has been reported to stop epileptic seizures [45]. The neurophysiological reason for the antiepileptic activity of random stimulation could be that the perturbation desynchronizes the cell networks that are progressively synchronizing during seizure development. Another common neurostimulation used to reduce seizures, vagal nerve stimulation (VNS), can cause a global change in excitability in several brain regions due to the vagus nerves anatomical connections, and the net effect is a decrease in neural synchronization that has been reported in patients that have shown reduced seizure frequency with VNS [46].

Furthermore, it is also of interest that neurostimulation methods that are nonspecific like electroconvulsive therapy (ECT), transcranial magnetic stimulation (TMS) and transcranial direct current stimulation (tDCS) are efficacious in the treatment of neuropsychiatric syndromes, perhaps due to the global nonspecific enhancement of brain excitability –the intensification of neural noise– achieved by these stimulation procedures. The specific cellular mechanisms by which the enhancement of variability in neural excitability is achieved varies depending on the method, but from a higher level perspective this phenomenon finds explanation in that an increase in the fluctuations in neuronal connections –or equivalently increasing the term *T*, “brain noise”, in the above formula– enhances the entropic term and helps reduce the free energy. It is thus tempting to speculate that a noisy brain –in terms of fluctuations in connections among brain regions– is a healthy brain, as has been asserted by some [47, 48].Whereas it is true that in some cases neurostimulation may promote inhibitory activity, sometimes this leads to excitation in other connected brain regions. Perhaps more accurate would be to say that neurostimulation alters the patterns of synchronous activity [49, 50]. Illustrations of the favourable effects on neuropathologies of nonspecific neurostimulation are numerous; to wit, there is evidence that DBS promotes memory [51] and non-invasive neurostimulation (rTMS and tDCS) enhances performances on several cognitive functions impaired in Alzheimer disease while some promising results with invasive DBS have also been observed [52]. In addition, non-invasive brain stimulation improves post-stroke recovery [53] and rehabilitation in general [54].

Why are these neurostimulation techniques able to improve brain healthy features? The reason, at a high level of description, may be the aforementioned tendency to increase dissipation, to promote the emergence and subsequent dissipation of energy gradients that increase the probability for neural connections to become active and engaged with the environment in efficient sensorimotor processing. These neural circuits may have already been moulded in the brain. The possibility of pre-configured neuronal functional connectivity motifs has been advanced by several scholars [55, 56], and the fact that there could be predefined dynamic patterns of neural activity and that external inputs re-activate what are already functional brain circuitries finds support in the abundant evidence obtained from invasive and noninvasive brain recordings indicating that there exist a common collection of network states [57, 58]. Even under anaesthesia correlations in brain activity have been found which are postulated to be not a reflection of sensorimotor processing –as there is very little under anaesthesia– but rather an intrinsic property of brain organization, perhaps helping to maintain or reinforce the already set connectivity patterns [59]. Repetitions of spontaneous patterns of neuronal activity, known as synfire chains, have even been found in vitro [60]. The spontaneous activity reverberating among these cell networks underlies the observations that the fluctuations of ongoing neuronal activity shape the stimulus-evoked responses [61].

Therefore, considering all these studies and the clinical data, the scenario that emerges is that of numerous possible functional connections among many neural networks with some of these remaining nearly continuously active. It almost looks as though in neuropathological states like coma, minimally conscious state or vegetative state some neuronal connections are not active but ready to become active, thus turning into the so-called functional connectivity which is based on their anatomical connectivity. Hence, neurostimulation procedures can make possible energy gradients that will make the connected cell ensembles functional again. But the crucial point for proper brain information processing, for conscious awareness to emerge, is that of fluctuating activity among many possible configurations of neural networks. Rather than stable states, brains need metastability for a correct function [13], whereas strong stability of brain states are normally associated with disease –e.g. status epilepticus, coma –or unconsciousness– slow wave sleep. In this regard, it has been shown that diverse patterns of correlated activity emerge from features that maximise variations in synchronization [62]. In the end, the notions of metastable brain states, variability in neural activity, long-range correlated activity and the need for widespread distribution of information for consciousness to arise, seem to have a common underlying theme: maximise the number of configurations of connections among neural networks, which from a high level description means the creation of energy gradients through efficient dissipation. Indeed, that fluctuations in functional connections among anatomically interconnected brain cell networks is a key to conscious awareness and healthy brain dynamics can be promoted to be a fundamental principle of brain function and organization. Therefore, and following the scheme presented in the first section, now the possibility of a probabilistic characterization of brain dynamics will be considered.

## 6 Probabilistic description of brain dynamics

We have seen how the entropy of the number of configurations of connected brain cell networks, associated with conscious awareness is higher than that during unconscious states and that the tendency of brain free energy is to decrease in the normal, wakeful states as opposed to pathological unconscious conditions. In other words, brain macrostates associated with conscious awareness possess more microstates (configurations of connections) whose emergence and dissolution determines cognitive states. The question now is whether the dynamical evolution of those microstates and the corresponding macrostates can be studied using an evolution equation. The usual procedure starts with a Langevin type of equation that encapsulates the deterministic and stochastic aspects. Here the proposal is to study the dynamics of the evolution of the brain coordination dynamics (synchrony patterns) starting from Shannons entropy, following the advice that “viewed in a dynamical perspective, generalised entropy-like quantities as used in information theory can provide useful characterizations of self-organising systems” [63]. In particular, it will introduce the perspective of the probabilities of interactions among neuronal networks as the basis to start developing an evolution equation.

The rationale for this probabilistic approach is founded in the collected works of several scholars. It has been said that If we are to understand biology we need a statistical mechanics of genes [64], which can be paraphrased as “If we are to understand the nervous system we need a statistical mechanics of nerve cell networks”. There have been efforts to characterise neurodynamics in terms of probabilistic notions, e.g. the early statistical neurodynamics [65], or the probabilistic descriptions frameworks [17], or [66] on the statistical mechanics of neocortex characterizing connections in terms of probabilities; all these ideas are reasonable because it is the global pattern of the many connections that is important and not the individual cell-cell connectivity. A host of computational models have been based on the probability of connections between units and have studied how these connections give rise to self-organised oscillations, e.g. [67]. It was mentioned in the introduction that correlations of activity (coherence, synchrony) can be considered a fundamental observable to describe brain states, therefore it is conceivable to use probabilities of functional connections –which will determine the correlated activity– among cell networks as a fundamental, elementary variable to study nervous system dynamics.

The starting point for a probabilistic perspective could plausibly be Shannon’s famous formula *S* = −Σ_*i*_ *p*_*i*_ ln(*p*_*i*_). One reason to choose this formalism is that it has been proposed that in the case of non-equilibrium systems, entropy (or information) in the Shannon sense is more adequate than classical entropy [68]. It is also true that whereas universal entropy principles have been discussed in several works [69] there is debate as to whether these principles can be formulated for nonequilibrium processes. Nonetheless, as aforementioned, the application of notions of equilibrium thermodynamics at the mesoscale level is feasible [7] and the careful interpretation of results may provide additional insights into the organization of the operations of the nervous system. We note as well that the Gibbs and the Shannon entropies, and indeed the thermodynamical entropy, are to a large extent equivalent [18, 70].

Hence, without further ado we start then with Shannons expression *S* = −Σ_*i*_ *p*_*i*_ ln(*p*_*i*_). To start addressing the dynamics one can differentiate the formula with respect to time

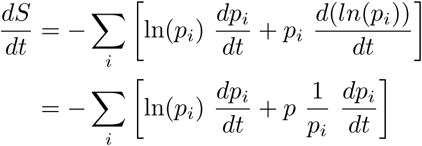

to obtain in the end

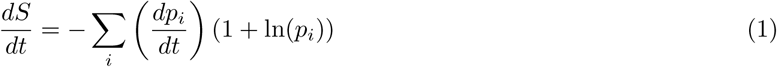

From the aforementioned rationale, *p* is the probability of functional interaction (or connection) between cell nets. The standard *p* in Shannon’s formalism refers to the probability of an event; in this case, the events are interactions between two cell ensembles. It could also be, as an alternative option, the probability of microstates– microstates in this case defined as aforesaid, combinations of pairwise connected networks – which, in the end, conceptually is the same as the previous notion: a probability of connection because microstates arise from connections. It is of interest that, recently, the thermodynamic entropy of a network (here networks defined as in graph theory) has been described using the Shannon formula dependent on the probabilities of the microstates [71].

From the above expression 1, one can immediately see that if *p* does not change (say, there is only one microstate, one configuration of pairwise connections that exists all the time), then there is no dissipation, *dS/dt* = 0, that is, the equilibrium state has been reached. In this case of *dp/dt* = 0 the steady state can be conceived as a pathological equilibrium, because fluctuations in connectivity patterns are crucial for proper brain information processing (sorry for belabouring the point!). It was noted above that dissipation increases with the magnitude of the fluctuations in connections, hence we arrive at the same result, but starting from another perspective, the Shannon formalism.

Assume now that *dp/dt* ≠ 0, then depending on the base of the logarithm there will be another steady state at which *dS/dt* is 0 when 1 + *ln*(*p*) = 0; the graph of *dS/dt* versus *p* is shown in Figure 5. Normally the logarithm is taken as the natural logarithm or at base 2, which suggests that the equilibrium *dS/dt* = 0 occurs around middle values of *p* = 0.36 in case of the natural logarithm or *p* = 0.5 if base 2 is chosen. Because, conceptually, the probability of connection is a function of the synchronization: *p* = *f* (*R*) (*R*, as described above, is the phase synchrony index, the higher it is, the more probable the connection and in fact *R* varies from 0 to 1 like the probability, so there could be a linear relationship between *p* and *R*), this qualitative account indicates that in normal brain function –this time the “healthy equilibrium”– the values of the synchronization to maintain a near equilibrium brain state should be in middle ranges, neither too low nor too high, which is what is normally found as opposed to high maintained values during coma or seizures (e.g., Figure 3). Other studies have presented evidence as well that consciousness requires medium values of certain features of neural assemblies [28, 72, 73].

**Figure 5:**
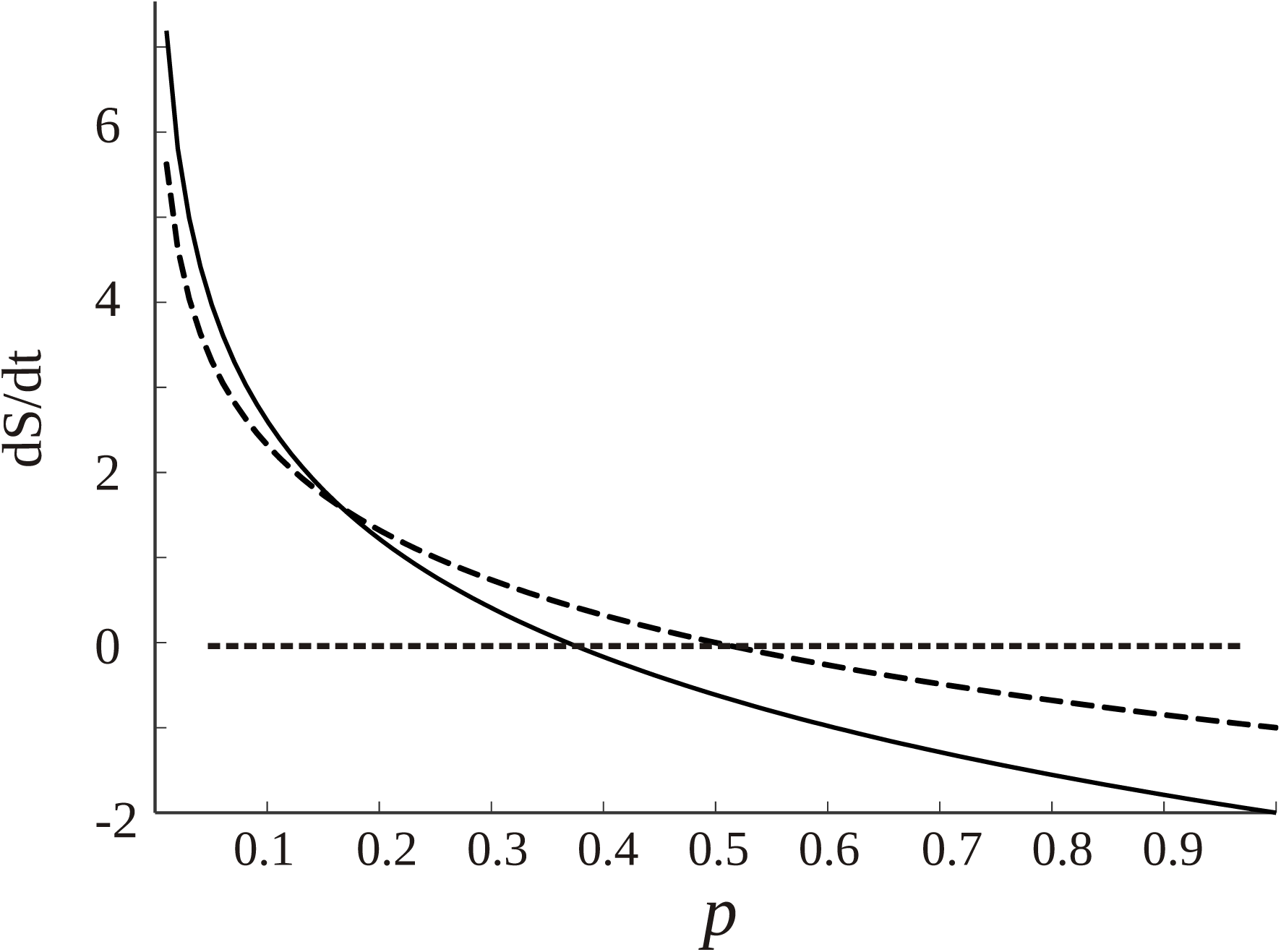
Dependence of *dS/dt* with the probability of connections p among cell networks, from equation 1 (assuming *dp/dt* = 1, if it has another constant value the graph is lifted following the y axis). The dotted curve is created using *log*_2_, and the black curve the natural logarithm. The straight dotted line denotes the equilibrium points at which *dS/dt* is 0. As commented in the text, there are healthy equilibrium points for middle ranges of *p*, depending on the base of the logarithm.

It has been emphasised that it is the instability, or metastability, of brain states that have adaptive values for the organism. Now we see that during normal cognition the state is near equilibrium, as it was seen as well in the graphs of the entropy of the macrostates (e.g. Figures 1 and 2 in Guevara Erra et al., 2016, current Figure 2), those during wakefulness being near the top of the Gaussian where equilibrium resides. This may be counterintuitive but the apparent paradox is resolved if one considers the global and local perspectives: the microstates (local view) need to fluctuate hence be unstable/metastable, whereas the global macrostate remains stable. This scenario of local instability and global stability is suggested by the inspection of the microscopic nature of the configurations of connections using a complexity measure derived from the Lempel-Ziv complexity –the normalised joint Lempel-Ziv complexity– applied to the connectivity among brain signals to evaluate the fluctuations in the connectivity pattern of the combination of networks in short time windows, which resulted in large values (that is, large variability) in wakefulness [30]. Thus, in those studies entropy and complexity measures were set in the context of connections among brain networks to evaluate the variability and the global information content of the system. In classical thermodynamics applied to chemistry the approach to equilibrium consists in a rearrangement of the mass distribution in a certain volume, but in the case of brains it is the arrangement of distribution of connected neural networks.

These conclusions of brain dynamics in conscious awareness being close to equilibrium are not inconsistent with the comments above about the higher dissipation during conscious alertness as opposed to unconscious states, because, as it has been many times mentioned, it is unfeasible to exactly quantify terms like *dS/dt*, and only qualitative tendencies can be discerned. In the brain macrostate associated with normal cognition near equilibrium there may be little dissipation low value of *dS/dt* yet more than during pathological states where due to the low fluctuations in synchrony the dissipation may be close to null. This is in fact seen in Figure 6. In order to inspect the changes in *dS/dt* in more detail, a simulation of equation 1 was performed using brain recordings from subjects previously analysed in [30]. Figure 6 depicts the time evolution of the dissipation in the transition from normal activity to an epileptic seizure. Note that there is an almost perfect flat line centred at 0 during the ictal events. To inspect the trends in *dS/dt* in another pathological condition, we computed the average values during states of coma as compared with the values in a healthy group, being 0.167 ± 0.04 (arbitrary units) for the former group and 0.196 ± 0.02 in the latter, hence again less dissipation is evident during this pathological unconscious state of coma. On the other hand, in a physiological (healthy) unconscious state, namely slow wave and REM sleep, there was no difference in the dissipation as compared to the awake conditions in the same individuals.

**Figure 6:**
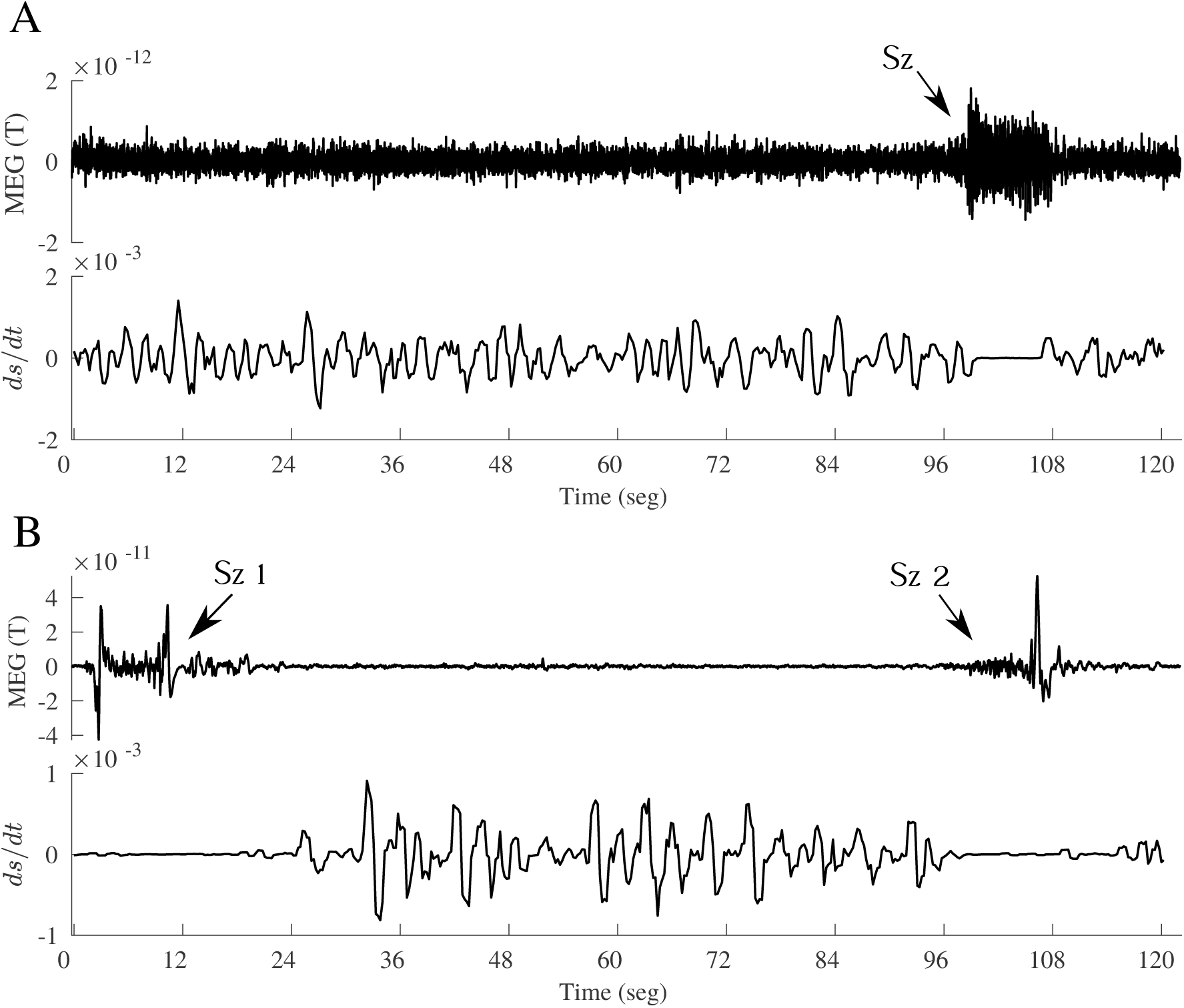
Time evolution of *dS/dt*, according to equation 1, in the transition from interictal to ictal events in two epileptic patients. **A**, upper panel shows a magnetoencephalographic (MEG) signal including an absence seizure (*Sz*) of about 10 sec duration, and the lower panel de evolution of *dS/dt* (arbitrary units). To compute this using eq. 1, the time evolution of the phase synchrony index *R* between two MEG signals (one shown above) was used, and the assumption was that *p* = *R*; then the *R* time series was divided into bins of certain duration from where the *< R >* was obtained for each bin and *dR/dt* evaluated as the slope from the first *R* value to the last value in each bin. This creates a *dS/dt* time series, which is plotted in the lower panels in A and B. **B**, another MEG signal, containing two seizures (*Sz*_1_ and *Sz*_2_), taken in a patient with frontal lobe epilepsy, demonstrating again the decreased dissipation (*dS/dt* close to 0) during and around the ictal events.

Taken together, these observations suggest that conscious awareness –that is, brain interacting with the environment– brings brain dynamics close to a “healthy” equilibrium with some, albeit small, dissipation needed to establish communication channels between cellnetworks. This situation in brain dynamics being close to equilibrium and of low dissipation may find its parallel in the case of biochemical oscillations; for instance, glycolytic oscillation in cells have been shown to be in near-equilibrium conditions with low dissipation so that small energy inputs can sustain the oscillatory, metabolic regimes [74]. These observations challenge the common view of metabolism being highly dissipative, and possibly this notion can be generalized to other biological processes operating in near-equilibrium regimes, like cognition.

The regions of positive and negative *dS/dt*, as seen in Figure 5, occur because of the simplifying assumption that *dp/dt* is constant. While positive and negative entropy production has been described in thermodynamic settings [75], here it is the Shannon formalism that is being applied and in this case the entropy is equivalent to the information content [76, 77]. Previous studies showed that the information content is larger in the network associated to conscious states [29] –perhaps indicating that consciousness could be the result of an optimization of information processing– and *dS/dt* would now be the variation of the information content. The region of positive variation of information with time suggests creation of information, occurring for low to moderate values of the probability of connections, whereas when the probability is higher, say in states like seizures, deep sleep or coma, there is decreased production of information. If generation of information can be attributed to healthy brain dynamics, the tendency thus is to have not too high synchronization (equivalent to high *p*); in other words, we find again the notion of variability in functional connectivity associated with healthy conscious awareness. We see that from different perspectives the same conclusion is reached.

But *dp/dt* may not be constant, in which case the graph of Figure 5 would change. In some works it has been taken as an operator representing probabilities to reach one state from another [63]. Another option would be to use a Fokker Planck equation (FPE): *dp/dt* = −*d*[*D*_1_(*x*) *p*]*/dx* + *d*^2^[*D*_2_(*X*) *p*]*/dx*^2^ with *D*_1_(*x*) and *D*_2_(*x*) the typical drift and diffusion coefficients associated with the FPE. Hence

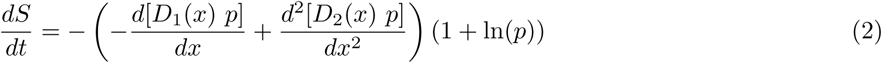

How to obtain the derivatives of p with the variable *x* ? What could be that variable? For example, *x* could be the phase synchrony index *R*, as it was abovementioned that *p* = *F* (*R*). It was as well noted that the synchrony index varies from 0 to 1 like the probability, so there could be a linear relationship between *p* and *R*: a maximum *R* (*R* = 1) indicates maximum probability of interaction (*p* = 1), and *R* = 0 indicates null functional connectivity, considerations that are more or less reasonable. For the sake for simplicity, assume a straight linear relationship. Therefore *dp/dR* = *K* (constant) and the second derivative is 0, so assuming *D*_1_ = 1 results in

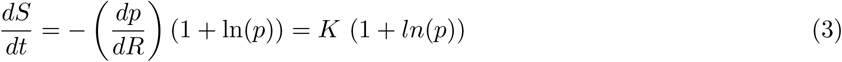

So with those assumptions the equation reduces to the previous 1 (with *dp/dt* = *constant*), hence same tendencies as described above will emerge from this FPE perspective (under the simplifying assumptions chosen here). These reflections, let us emphasise again, are not meant to quantify precisely these expression but rather to inspect qualitative tendencies.

## 7 The emergence of cognition from the previous elaborated perspective: compartmentalization plus dissipation

The main purpose, as mentioned at the beginning, is the search for a simple unified conceptual framework to describe the organization of brain dynamics. Can these results obtained here be bound together into some relatively simple coherence with a central theme? The main findings all revolve around the notion that the brain macrostates comprised of neural networks connections associated with conscious awareness and health had more configurations –more microstates– thus conferring brain areas more variability to establish different connectivity patterns for proper sensorimotor transformations, reminiscent of old proposals like Flohrs rate of dissolution of neural assembles determining degrees of consciousness [78]. It may be worth now to take a look at what biology teaches us, for after all cognition is an aspect of it.

Compartmentalization is considered the first step in biological organization [79], that joined with dissipation gives rise to self-organization, the formation of complex structures: “Self-assembly and a process of dissipative structure formation are complementary principles of self-organization” [80]. Can therefore compartmentalization plus dissipation provide the basis for conscious awareness?

In biological systems compartmentalization is normally achieved by aggregation of lipids, forming micelles. At higher levels, modules of connected cells appear. The emergence of modularity has been studied for long time and is considered to represent an advantageous organization for organisms [81]. In the brain a compartment can be thought of as a number of cell networks that are synchronous, or functionally connected. Indeed, the modular (in anatomical sense) construction of invertebrate and vertebrate nervous systems has been known for long [82]. Anatomical and neurophysiological studies have taught us that nervous systems are organised in a modular fashion, e.g., neuronal tract tracing unveiled modular organization consisting of connections linking regularly spaced cell clusters [83, 84]. The modular organization in brains may underlie the common motifs observed in EEG and other brain recordings that were mentioned above in the section on improving healthy brain dynamics [57, 58].

It is worth noting some similarities in the formation of brain cell functional clusters with the formation of micelles. This is a highly cooperative process that depends on the lipid (monomer) concentration. In the case of nervous systems, the synchronization between two networks is also cooperative and depends on how many cells are active, the more neurons firing in synchrony the more probable the connected networks will start becoming active and the stronger the electric field will be around the neurons that, in turn, will enhance activity –the so-called ephaptic coupling [85]; this phenomenon is related to the enslaving principle of synergetics, where the field enslaves the units. The size of micelles is determined by geometric and thermodynamics factors, and in the case of neuronal functional connections these are determined as well by the geometry of the anatomical connectivity, whereas the thermodynamic factors favouring synchronization can be those expounded in previous sections: the increase in dissipation and reduction of energy gradients and free energy. The advantage for those investigating membrane formation by lipids is that the thermodynamic drive can be quantified by means of a very useful notion, the chemical potential: the organised structures are formed by the molecules searching for their lowest potential. But there is no equivalent to a chemical potential for the collective activity of brain cell ensembles. Could there be an analogous notion that applies to brain function and cognition? A subsection below addresses this topic.

A main difference between the biological compartments and those compartments in the brain formed by the neuronal functional connections is that the former tend to be stable while the latter are transient (except in pathological and unconscious states). While this partition into cell clusters proposed here is considered at the neurophysiological level, other more abstract partitions such as in mental state space have been advanced with the proposal that stable partitions of microstates give rise to cognitive macrostates that depend on context and environment [86].

Perhaps then, compartmentalization, the transient formation of neuronal connections, may be one fundamental aspect that is entwined with the other aspect, energy dissipation that creates and destroys the functional cell assemblies. In attempts to explain the emergence of complexity, it has been advanced that the mechanisms of systems moving down energy gradients is what leads to complexity (Schneider and Sagan, 2005), this being related to Prigogines proposal that dissipation of energy creates organization. Conceivably, the emergence of complex structures requires many interacting units exchanging energy, and this could be the mechanism how energy moving down gradients creates complexity, by maximising the number of interactions among units that exchange that energy. According to the aforementioned results, in brain cell circuitries the energy gradients that establish functional cell connections maximise the configurations of connections during conscious awareness, maximisation of units (cells) exchanging energy –or information, if you will. It is tempting to conjecture that because brains function mainly to maintain a model of the environment, then perchance the brains global dynamical structure has to copy what is out there, the dynamical structure of the world which consists in energy being dissipated and distributing in all possible microstates, according to the second principle of thermodynamics.

In closing, it is of interest that already in the early 1900s A. J. Lotka proposed that “natural selection [and evolution] tends to make the energy flux through the system a maximum, so far as compatible with the constraints…” [87]. It can be thus envisaged that biological (consciousness) evolution occurs as to make the energy flux through the system maximum among all constituents (brain cells) compatible with constraints from the environment. The expression of this feature in the function of the nervous system and especially the brain could be the main factor accountable for the emergence of conscious awareness. This maximisation of energy spread may account for the many times proposed maximisation of entropy or information. But some scholars are reluctant to consider these maximisation principles as cause of processes driving living systems [68]. It may be so, because concepts like information or entropy are, after all, our creations that were developed to characterise certain phenomena. It is worth noting that Haken’s perspective emphasises tendencies in natural phenomena –specifically about the emergence of macroscopic order– rather than precise quantifications. This focusing on the search for tendencies is a view that, as many times abovementioned, has been taken through this text, and it is perhaps the tendency towards minimization of free energy favouring the spread of energy –that becomes manifest through interaction among the system’s constituents exchanging energy– which underlies the said principles about entropy and information. This notion makes the energy perspective a most fundamental aspect to understand not only cognition but the organization of natural phenomena in general [14].

### 7.1 A brief digression on possible foundations for a conceivable nervous system potential

It was mentioned above the usefulness of having a chemical potential to characterise the evolution of chemical reactions. J.W. Gibbs definition of chemical potential in 1875 allowed for the precise characterization of aspects like equilibrium and the investigation of the dynamics of chemical reactions. It could be of interest if some equivalent to the chemical potential could be described about the processes of the nervous system –especially in its collective cellular activity at the meso/macroscale level– a concept that could help understand the evolution of brain activity and its relation to behavioural dispositions. It was described above that in the expression for the entropy production *dS/dt* the term *N*_*i*_ could be taken as pairwise connected brain networks instead of molecules. Hence, the basic intuition behind the idea to develop an analogue to chemical potential for the brain is that connected cell ensembles are analogous to molecular concentrations in chemistry.

The chemical potential, *µ* of a component in a solution can be thought of in several ways, as a measure of the “escaping tendency” for a component in a solution and as a measure of the reactivity of a component. By analogy, the “nervous system potential” (lets call it *µ*_*NS*_) can be thought of as a measure of the formation and dissolution (escape) of functional cell assemblies and as a measure of the connectivity tendencies, the ‘reactivity’ of cells establishing connections. The chemical potential refers to ‘components’ in a (solid, liquid or gaseous) ‘solution’, the *µ*_*N*_ *S* refers to ‘cell assemblies’ in the nervous system ‘tissue’. In this manner, the main building blocks of brain activity at mesoscale levels are cell assemblies functionally connected, which is a current concept in neuroscience and was described in the introductory paragraphs. The dynamic evolution of chemical systems is well established: molecules “search” for their lowest *µ*, and in equilibrium there is no further change in the number of molecules, just fluctuations around the equilibrium state. In our studies we have observed that the entropy associated with states of conscious awareness are near the equilibrium; this indicates that the number of connected cell networks remain stable but there are fluctuations, and it is the fluctuations in connectivity that, as already emphasised above, is a crucial factor for proper brain information processing.

Nevertheless the parallel between *µ* and *µ*_*NS*_ has difficulties when it comes to measuring “concentrations” of cell assemblies, and as a result the standard derivation of the expression for *µ* in a chemical reaction system cannot be achieved for *µ*_*NS*_ because it starts from the Gibbs free energy formula whose terms have clear meaning when applied to chemistry. For each molecular component *i*, the chemical potential is defined as 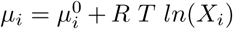 where *X*_*i*_ is the molar fraction, which may not have and easy parallel in the case of the brain cell networks. Neither *µ*_*i*_ has easy interpretation in case of brain neuronal networks. The energy (enthalpy) term does not appear in the formula due to considerations of mixing several molecular entities, considerations which would not apply to brains.

Now, one option that may have promise would be to define *µ*_*NS*_ for each macrostate, a macrostate being the number of configurations of connected cell networks as was done in our estimation of entropies. In this case while the entropy and the fluctuations in connections can be quantified for each macrostate, the problem of assigning an energy still remains, as commented above. Nonetheless, the expression would be then for brain macrostate *i, µ*_*i*_ = *E*_*i*_(*dN*_*i*_*/dt*) *S*_*i*_ where *S*_*i*_ is the entropy of the macrostate and *dN*_*i*_*/dt* represents the “temperature”, or fluctuations in the connections of the network components of that macrostate that can be evaluated over a certain time interval.

## 8 Conclusions

As a manner of closing these arguments that try to shed light into brain organization in cognition, and to make them applicable to the understanding of some neuropathological syndromes, the following scheme (Figure 7) depicting the scenario towards the transition to brain pathological activity can be envisaged [88]. We also briefly note that the maximisation of configurations of synchrony and the associated dissipation of gradients concurrent with energy fluxes that help organise the system providing largest number of metastable states is very related to the notion of criticality in nervous system dynamics [89], because near equilibrium where there is large number of configurations of cell network connections allowing to handle proper information processing in the brain, the system is highly susceptible to inputs and therefore fosters adaptability of the organism. The key then is not to reach the maximum number of units interacting (which would be all-to-all connections and thus only one possible microstate), but rather the largest possible number of configurations allowed by the constraints, in other words, a critical state. Departure from criticality has been noted in unconscious states [90].

**Figure 7:**
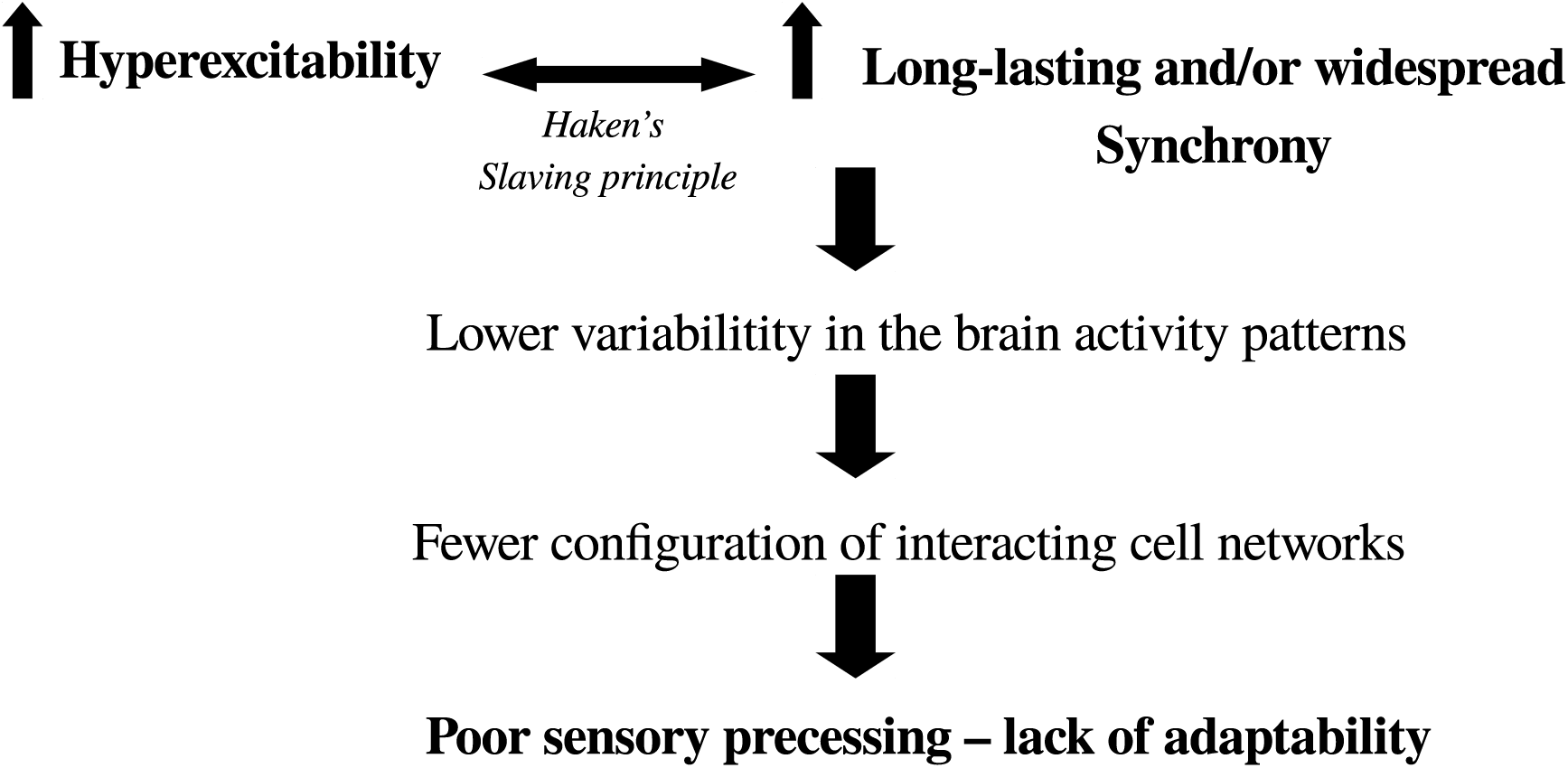
Diagram depicting the scenario towards the transition from normal to abnormal brain activity. It is based on two main features: hyperexcitability and higher-than-normal synchrony. Where in the brain these two events occur will determine the pathology (e.g., in Parkinson’s there is increased excitability and synchrony in basal ganglia-thalamocortical networks). It is well-known that enhanced cell firing leads to pronounced synchrony which, if it is too stable, results in low uctuations in activity leading to fewer combinations of functional connectivity (lower complexity, departure from criticality) and thus the brain has diffculty processing sensorimotor transformations in a fast and variable manner to guide normal, adaptive behaviour.

A direct test for the framework proposed in this text would be the demonstration that optimality of sensorimotor processing is associated with near maximal number of possible configurations of brain networks –higher entropy of the neuronal connectivity– and with enhanced short-time scale fluctuations in the connectivity among those networks. While it was previously observed that lower entropy was found when visual input was diminished –subjects with eyes closed ([29]) and as well similar results of more regular, less complex, network structure as compared with eyes open condition in [91]– and that lower fluctuations (given by a complexity measure) of connectivity was associated with epilepsy and coma, there are many other situations and pathological conditions where this assessment could be performed, like vegetative or minimally conscious states. But still remains to find an adequate evolution equation for the general principle proposed here, from which specific outcomes may be put to the test. The present work has been an attempt to start considering this from a high-level perspective that, we posit, may yield a comprehension of the brain∼behaviour relation.

